# Rapid bilevel optimization to concurrently solve musculoskeletal scaling, marker registration, and inverse kinematic problems for human motion reconstruction

**DOI:** 10.1101/2022.08.22.504896

**Authors:** Keenon Werling, Michael Raitor, Jon Stingel, Jennifer L. Hicks, Steve Collins, Scott L. Delp, C. Karen Liu

## Abstract

Creating large-scale public datasets of human motion biomechanics could unlock data-driven breakthroughs in our understanding of human motion, neuromuscular diseases, and assistive devices. However, the manual effort currently required to process motion capture data is costly and limits the collection and sharing of large-scale biomechanical datasets. We present a method to automate and standardize motion capture data processing: bilevel optimization that is able to scale the body segments of a musculoskeletal model, register the locations of optical markers placed on an experimental subject to the markers on a musculoskeletal model, and compute body segment kinematics given trajectories of experimental markers during a motion. The optimization requires less than five minutes of computation to process a subject’s motion capture data, compared with about one day of manual work for a human expert. On a sample of 34 trials of experimental data, the root-mean-square marker reconstruction error (RMSE) was 1.38 cm, approximately 40% lower than the 2.58 cm achieved manually by 3 experts. Optimization solutions reconstructed known joint angle trajectories from four diverse motion trials of synthetic data to an average of 0.79 degrees RMSE. We have published an open source cloud service at AddBiomechanics.org to process experimental motion capture data, which is available at no cost and asks that users agree to share processed and de-identified data with the community. Reducing the barriers to processing and sharing high-quality human motion biomechanics data will enable more people to engage in state-of-the-art biomechanical analysis in their work, do so at lower cost, and share larger and more accurate datasets.

**Author summary:** Creating large-scale public datasets of human motion could unlock data-driven breakthroughs in our understanding of neuromuscular diseases, assistive devices, and human motion more broadly. The manual effort currently required to process these motion datasets is costly and limits the collection and sharing of large-scale datasets. Our cloud-based software tool, called AddBiomechanics, uses state-of-the-art optimization techniques to automatically scale the body segments of a musculoskeletal model to match the subject of interest, and then compute body segment kinematics during a motion. The optimization requires less than five minutes of computation to process a subject’s motion capture data, compared with about one day of manual work for a human expert. The accuracy of the approach in quantifying the body segment kinematics is as good or better than the results achieved manually by experts. Reducing the barriers to processing and sharing high-quality human motion biomechanics data will enable more people to engage in state-of-the-art biomechanical analysis, do so at lower cost, and share larger and more accurate datasets.

## Introduction

Quantitative analysis of human movement dynamics is a powerful tool that has been widely used to solve problems such as assessing muscle and joint function e.g. [1–5], better understanding joint loading and consequent adverse health outcomes e.g. [6–12], analyzing the performance of assistive devices for improving human movement e.g. [13–16], quantifying the effects of complex conditions like Parkinson’s disease e.g. [17,18], and even generating more realistic computer graphics e.g. [19,20]. The benchmark data acquisition technique for quantifying human biomechanics remains optical motion capture [21, 22]. This process involves placing a number of optical markers on a subject’s body segments and having the subject perform actions in a laboratory space surrounded by specialized cameras. These camera systems are able to reconstruct the three-dimensional locations of the optical markers in the lab, and given the marker trajectories over time, one can use proprietary, open, or custom software to reconstruct the kinematics of the subject’s body segments.

The current state-of-the-art software for reconstructing the motion of a human musculoskeletal model from optical marker trajectories requires substantial iterative “guess-and-check” refinement, and can take a large amount of expert time to achieve a high quality result on each new subject. This manual interaction increases costs, limits scalability, and reduces the reproducibility of motion capture studies [23,24].

To reconstruct movement kinematics from optical motion capture data, software must address several sources of noise, ambiguity, and model error. Initially the optical markers all appear as undifferentiated bright spots to the cameras, so the first challenge is identifying which bright spots correspond to which locations of markers placed on the body. This is called *marker labeling,* and is generally handled by having a user manually label each marker using commercial software from motion capture vendors [25, 26]. Once the markers have been labeled, software must reconstruct a digital twin of the captured subject, with segment dimensions that match the physical subject as closely as possible. This process is called *scaling,* and a variety of approaches have been described [23,27–34]. Finding accurate scaling is especially important when using motion capture data to create muscle-driven simulations because the muscle-tendon parameters are scaled by the body segment dimensions [35]. To achieve accurate kinematic results, the locations of the markers on the scaled digital twin must be adjusted to account for variations caused by human error in attaching the markers to the body and the variations in the dimensions of human subjects [24]. This is called *marker registration.* Finally, the positions and orientations of the body segments over time must be determined, which is typically done using an optimization process called inverse kinematics [36–40]. Inverse kinematics algorithms generally produce more accurate results when the solutions are constrained by an underlying skeletal model [14, 24, 41].

In practice, the interdependence between scaling, marker registration, and inverse kinematics means that experts must follow an iterative guess-and-check procedure, where they refine each of the steps several times, making small adjustments to each value until a desired accuracy is achieved [42,43]. For example, increasing the length of the upper arm segment in a subject’s digital twin will require also changing the marker registrations for any markers on the forearm and the hands, because otherwise those markers would move as a result of the longer upper arm. A longer upper arm will also, all else being equal, change the resulting motion found by inverse kinematics. While there are best practices for conducting validation at each step [35], the process typically requires subjective input from an expert and time-consuming interaction.

Automating the scaling and registration process has been studied before, in the pioneering work of Reinbolt et. al. [44] and Charlton et. al. [45]. These papers found it is possible to use gradient-free optimization methods to automatically guess body segment scales and marker registrations while solving gradient-based inverse-kinematics problems repeatedly in an inner-loop to evaluate optimization progress. These methods require large amounts of compute time because every iterative guess the outer optimizer makes about body segment scaling and marker offset must solve its own inner optimization problem (inverse kinematics) to evaluate the quality of the guess, which is itself computationally costly. The method of Reinbolt and colleagues [44] also finds best results using a particle-based optimizer for their outer optimization problem, to combat the non-convexity of the problem, but this comes at a further multiple of computational cost.

Given the interconnected nature of body segment scaling, marker registration, and inverse kinematics, one might also consider posing all three problems as a single optimization problem. However, such a formulation leads to a nonconvex optimization in which a global solution is not guaranteed [46]. Instead, we can only guarantee to find a local optimum close to an initial guess, so providing a high quality initial guess for parameters is crucial. Andersen and colleagues [47] have formulated such nonconvex optimization problems, but did not address the problem of reliably finding an initial guess for the non-convex optimization problem proposed.

This paper introduces an automated method (Fig 1) that uses a combination of traditional kinematic solvers and modern bilevel optimization techniques to achieve high quality results in reasonable computation time. Our method requires neither a user-provided initial guess for the decision variables nor large computational resources. We first apply a sequence of simple optimizations to individually approximate the initial values for the body segment scales, marker registrations, and inverse kinematics [44, 45]. Then, rather than iteratively repeat those optimization problems hundreds of times as in previous work, we apply bilevel optimization techniques to simultaneously optimize body scaling, marker registration, and inverse kinematics. This process takes approximately 3-5 minutes on a consumer grade laptop computer or low-end server.

**Fig 1.**
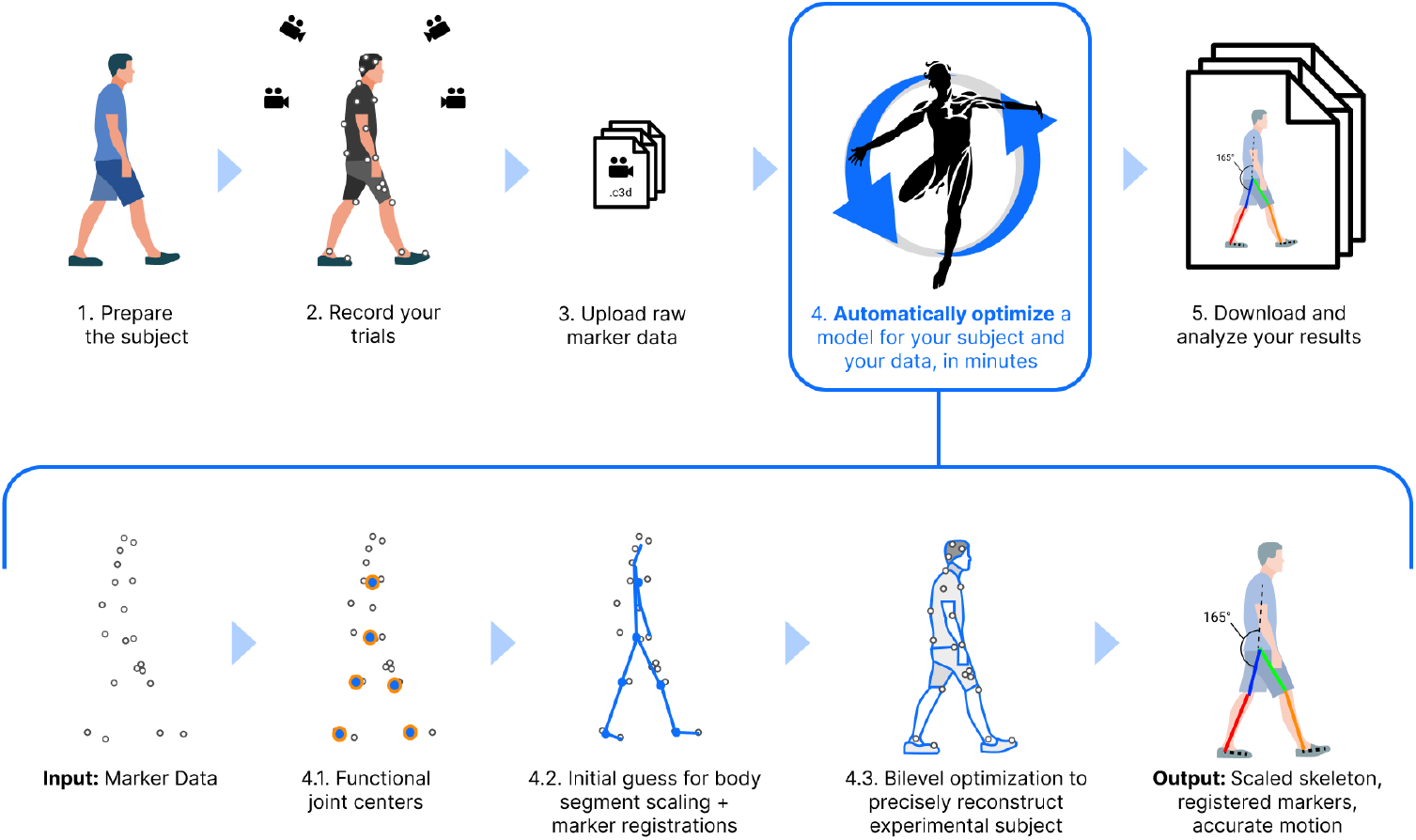
How our method fits into an optical motion capture pipeline. AddBiomechanics integrates into the standard optical motion capture pipeline to automate the process of model scaling, marker registration, and inverse kinematics. Once raw marker data has been collected and uploaded (steps 1-3), addBiomechanics (step 4), replaces the time-consuming and error-prone manual step in previous workflows. Our method processes input marker data through several steps automatically. First, it finds the functional joint centers from the data (step 4.1), and then it uses the marker data and those joint centers to make an initial guess for body segment scaling and marker registrations (step 4.2). The initial guess then serves as the starting point for a bilevel optimization problem that matches the model to the experimental data as closely as possible (step 4.3). The final output is a musculoskeletal model scaled to the subject with registered markers and joint angles over time.

Our approach is appropriate for use by non-experts to process large amounts of motion capture data automatically. We have released the software, which we call “AddBiomechanics,” as open source. We also provide a cloud-based service hosting AddBiomechanics, available at addbiomechanics.org, where researchers can process data without downloading or installing any software. We provide a drag-and-drop interface that has the potential to save thousands of hours and enable quantitative movement analysis without arduous manual steps, while simultaneously improving the quality of results. AddBiomechanics outputs OpenSim project files [42], compatible with the widely used open source biomechanics package, so the results of scaling and marker registration can be transferred to OpenSim for further analysis.

## Methods

Given the measured marker trajectories of length *T*, 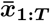 from a motion capture system, concurrently reasoning about model parameters and data is formulated as a nonconvex optimization that solves for the kinematic pose trajectories, **q**_1:*T*_, the scaling parameters of the body segments of the musculoskeletal model, ***s***, and the locations of markers attached to the body segments, ***p***. The objective of the optimization is to minimize the deviation of estimated marker positions from 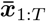:

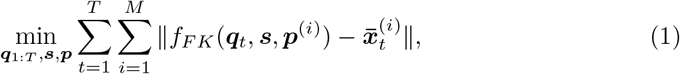

where *M* is the number of markers, and *f_FK_* (***q**_t_*, ***s***, ***p***^(*i*)^) is the forward kinematic process that transforms a point ***p***^(*i*)^ on the body segment for marker *i* in a skeleton scaled by ***s*** in the pose ***q**_t_* from the local coordinate frame of the assigned body segment to the world coordinate frame. ***p***^(*i*)^ ∈ ℝ^3^ denotes the local position of the i-th marker while ***p*** ∈ ℝ^3×*M*^ is the concatenated local positions of all markers.

Equation 1 is high-dimensional and nonconvex. Consequently, the solution of such an optimization is highly sensitive to the initialization of the decision variables. We introduce a bilevel maximum-a-posteriori (MAP) optimization and an initialization strategy to achieve new state-of-the-art in automatic processing of biomechanical motion capture data. The proposed bilevel MAP optimization simultaneously considers data reconstruction and anthropometric statistics when jointly optimizing all decision variables in Equation 1. To overcome the sensitivity to the initial guess, our method individually initializes each type of variable utilizing independent sources of information. Specifically, we use kinematic constraints to initialize ***q***_1:*T*_, a geometric invariant to initialize ***s***, and real-world measurement to initialize ***p***. Once the variables are initialized individually, the final bilevel optimization simply makes sure that they agree with one another, given the observed data and model priors.

### Input Marker Data

The output of a commercial motion capture system is a series of frames, often at 100-200 Hz, where each frame contains a variable number of 3D coordinates representing the locations of optical motion capture markers in the experimental capture volume at the moment in time corresponding to this frame. Each 3D coordinate is “labeled” with a tag corresponding to an experimental marker location on the subject (e.g. “C7” for the optical marker placed on the C7 spinal segment). A full list of marker tags, and their location on an idealized digital twin of the experimental subject, is known as the “marker set.” In practice, markers are almost never placed *exactly* at their ideal locations, and these small deviations in experimental marker placement must be accounted for during the marker registration step. Not every marker from the marker set is observed on every frame, because they may be occasionally obstructed during a motion capture experiment. The only assumption we make is that the tag associated with an observed marker is correct.

### Input Unscaled Skeleton

Our algorithm can accept arbitrary skeletons in OpenSim format to scale and register. A skeleton is parameterized by a set of body segments, connected by joints. The scaling of each link is concatenated to form the ***s*** vector, and the degrees of freedom of each joint are concatenated to form the ***q*** vector. The algorithm supports all OpenSim joint types, including custom joints. Examples of skeletons that have been successfully scaled and registered in our experiments include widely used state-of-the-art biomechanical models [48,49].

### Bilevel Maximum-a-Posteriori (MAP) Optimization

The manual method for processing motion capture data requires that the expert perform the following steps in a loop: manually adjust the body scalings and marker offsets, run inverse-kinematics (IK) using the new parameters, visually inspect for errors, and repeat. The expert is engaged in a form of optimization problem (“find the best body scalings and marker offsets”) which involves calling a smaller optimization problem in the inner loop (running IK). We can mirror this nested optimization objective, where evaluating the objective function at a particular time requires running an optimization problem with bilevel optimization.

Given recorded marker positions 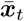 at time instance *t*, we are interested in reconstructing the scales of each body segment in the musculoskeletal model, ***s***, the local positions of the markers ***p*** attached to their assigned body segments, as well as the joint pose ***q**_t_*. This problem can be formulated as a maximum a-priori (MAP) optimization:

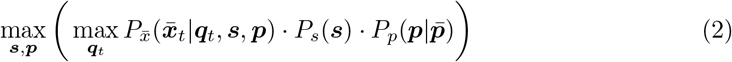

The first term is a conditional probability of the observed data given the estimated parameters. This formulation is equivalent to the standard least-squares inverse kinematics objective term if we assume Gaussian noise in our optical marker observations. The second term expresses the prior of skeleton scaling, encoded as a multivariate Gaussian fit to the ANSUR II dataset [50] of anthropometric scalings. If the height, weight, or biological sex of the experimental subject is known, the multivariate Gaussian skeleton scaling prior is conditioned on that information before any optimization. The third term is a zero-mean Gaussian distribution that regularizes the deviation of the marker locations from their intended locations 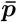 provided by the experimenter, encoding that markers are generally placed close to their intended locations, even if they do not perfectly align.

Note also that this is a bilevel optimization problem, because in order to evaluate the quality of given skeleton scaling ***s*** and marker locations ***p***, we need to optimize over the possible joint positions ***q_t_***. To efficiently solve the bilevel optimization problem, we observe that at the optimal values of ***q**_t_* for max_*q_t_*_ 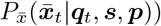, the gradient of the inner optimization problem will be zero. Using this observation, we reformulate the bilevel optimization problem as a single-level nonconvex optimization problem with nonconvex constraints. For numerical stability, we minimize the negative log of the above objective function:

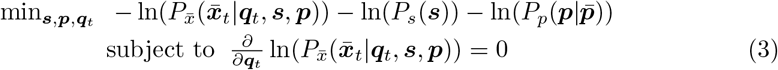

At a locally-optimal point, the gradient of the objective term with respect to any of the decision variables is zero, so it must be zero with respect to ***q**_t_*:

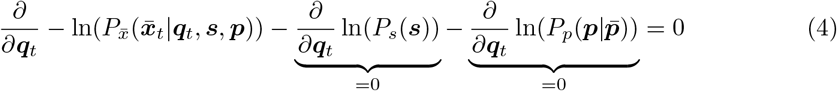

Thus, at a locally-optimal point for the objective function our constraint must hold regardless, and so we could theoretically omit it from the optimization problem without loss of correctness. However, we found that including the constraint explicitly allows the optimizer to converge to a high-quality solution much more quickly.

We could use any nonlinear optimization solver to solve Equation 3. In practice, we use IPOPT [51], which is a high-quality and open source solver. However, due to our problem’s non-convexity, a good initial guess for the decision variables is needed to produce reasonable results.

### Initializing Decision Variables

Prior to solving the optimization problem in Equation 3, we need to get “close-enough” initial guesses for the decision variables. We do this through a sequence of optimization problems shown as a flow-chart in Fig 2. We obtain initial guesses for ***q**_t_*, ***s***, and ***p*** individually based on independent sources of information such that the cascading errors can be mitigated.

1. Initialize ***p*** using the marker locations measured by the experimenter or defined by the existing marker set.
2. Initialize ***q**_t_* by solving inverse kinematics with a generic unscaled skeleton and marker locations provided by the previous step.
3. Initialize ***s*** by computing the functional joint axis and scaling ***s*** to match the joint axis along with the measured markers.

**Fig 2.**
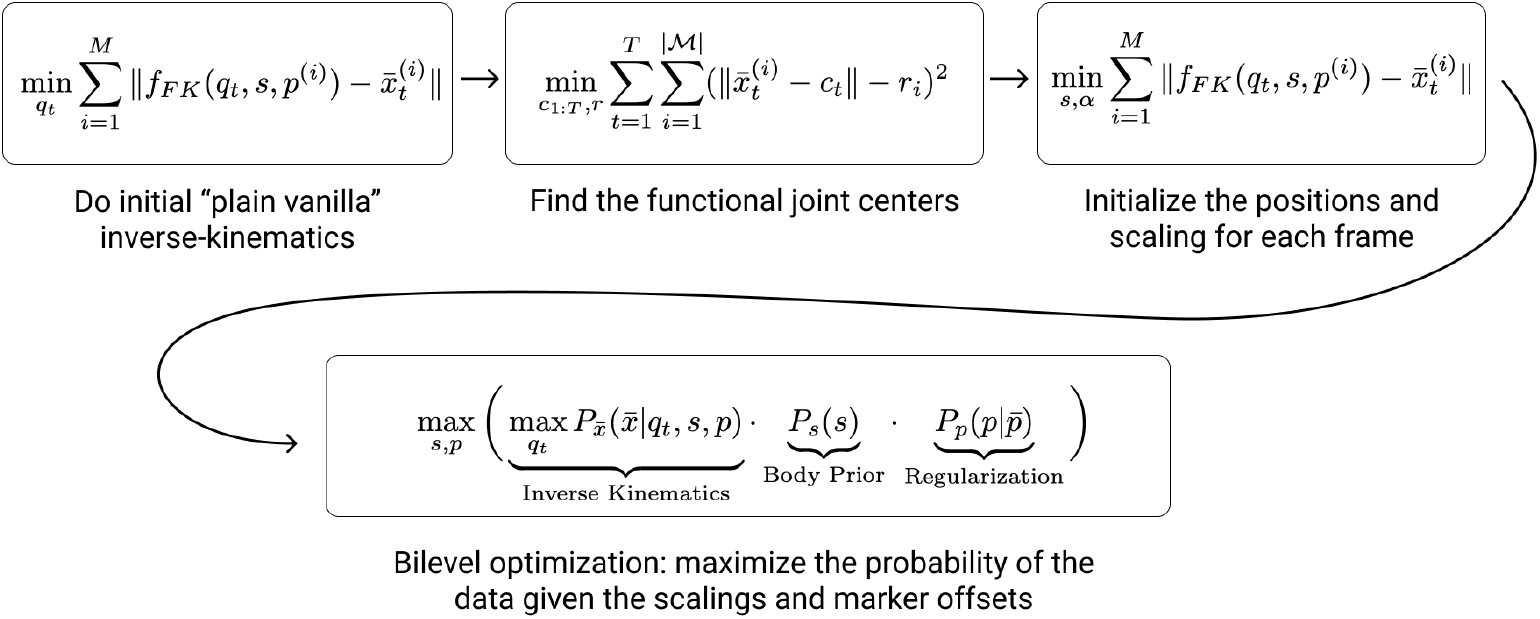
Initializing Decision Variables. This flow chart details the sequence of non-convex optimization problems that we solve. Each problem provides output that is useful for initializing the next non-convex optimization problem, steadily increasing in complexity.

Step 1 is trivial and Step 2 is a simplified Equation 1 with ***s*** being the population average and ***p*** given from Step 1, 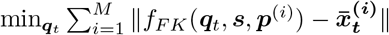. Solving this IK problem efficiently has been an area of research for decades [52–54] and can be done efficiently and reliably.

The most involved step in our initialization process is Step 3, initializing the body segment scales ***s***. We begin with computing a functional joint center from the measured marker trajectories. Let the subset of markers attached to the two bone segments connected by the joint be 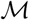, we can estimate the joint position ***c*** in the world frame over time by

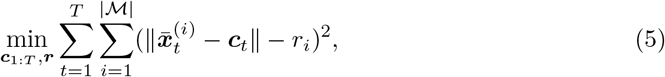

where *r_i_* is the estimated distance between the *i*-th marker 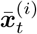 to the joint center ***c**_t_* for all *t*. *r_i_* is constant over time. For each marker, Equation 5 fits a moving sphere centered at ***c**_t_* with the radius *r_i_*, to match the measured positions of the marker over time.

The sphere-fitting approach to finding functional joint centers can yield ambiguities when marker motion adjacent to a joint is primarily confined to the sagittal plane, as commonly happens in locomotion. In such cases, we could move our joint center perpendicular to the sagittal plane, and still have an equally good solution for sphere-fitting. As a result, it might incorrectly scale the skeleton to match erroneous joint positions. For example, it might incorrectly scale the hip width while still matching all the measured marker motion for the thighs and the pelvis.

To address these ambiguities, we formulate another optimization problem to simultaneously find the joint axis and the joint center, building on Equation 5. This problem is similar in spirit to the axis-of-rotation problem described in [55], but can be implemented without any matrix factorizations. The goal of the axis fit problem is to identify not only a joint center ***c***, but also the direction of axis ***a*** at each frame. We also estimate a fixed distance from the center for each marker, parameterized by a distance *u_i_* along the axis ***a*** and a distance *v_i_* perpendicular to the axis ***a***. The result of a successful axis fit is that we capture a line at each frame, where the functional joint center could lie anywhere on that line:

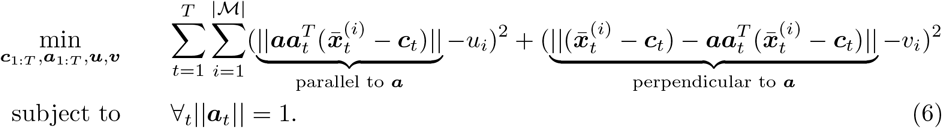

For each marker 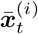 in the set 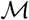, we decompose 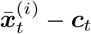 to two vectors: the parallel vector which is the projection of 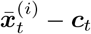 on ***a_t_***, and the remaining orthogonal vector. Equation 6 encourages that both the projected vector and the orthogonal vector maintain constant length over time for every marker in 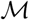.

We run both sphere fitting and axis fitting at each joint. Because the axis fit is a strictly more demanding problem, if we succeed in axis fitting, then we pass the axis on as a constraint for subsequent problems. If we fail at axis fitting, then it must be because there is out-of-plane marker motion, which means that our sphere fit is not ambiguous, so then we pass along the exact joint center to subsequent problems.

Since Equation 5 and 6 are nonconvex, we initialize ***c***_1:*T*_ using the joint trajectory ***q***_1:*T*_ solved by Step 2 and use any nonconvex optimization solver to optimize them. Once we determine the joint center and the joint axis, we formulate another optimization to initialize the scaling parameters ***s***:

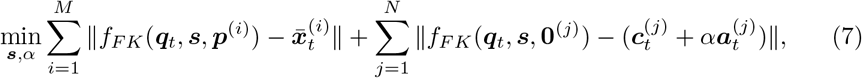

where the zero vector **0**^(*j*)^ indicates the local coordinate of the joint *j* and *N* is the number of joints. The first term fits the skeleton to the measured marker positions, while the second term encourages the joints to lie on the estimated joint axes solved by Equation 6, at a distance controlled by the scalar decision variable *α*. If the joint axis does not exist for the joint *j*, we set ***a**_j_* to zero and remove *α* from the optimization.

### Open Source Implementation

To facilitate adoption, we provide the algorithm as an open source cloud-based tool that allows researchers to automate scaling and marker registration on their motion capture data without downloading or installing any software, available at addbiomechanics.org. Users can drag and drop files for automated processing, and then visualize on the web or download results for analysis in OpenSim (Figure 3). The cloud tool also allows researchers to automatically generate comparisons of their own hand-scaled data versus the output of the automated system.

**Fig 3.**
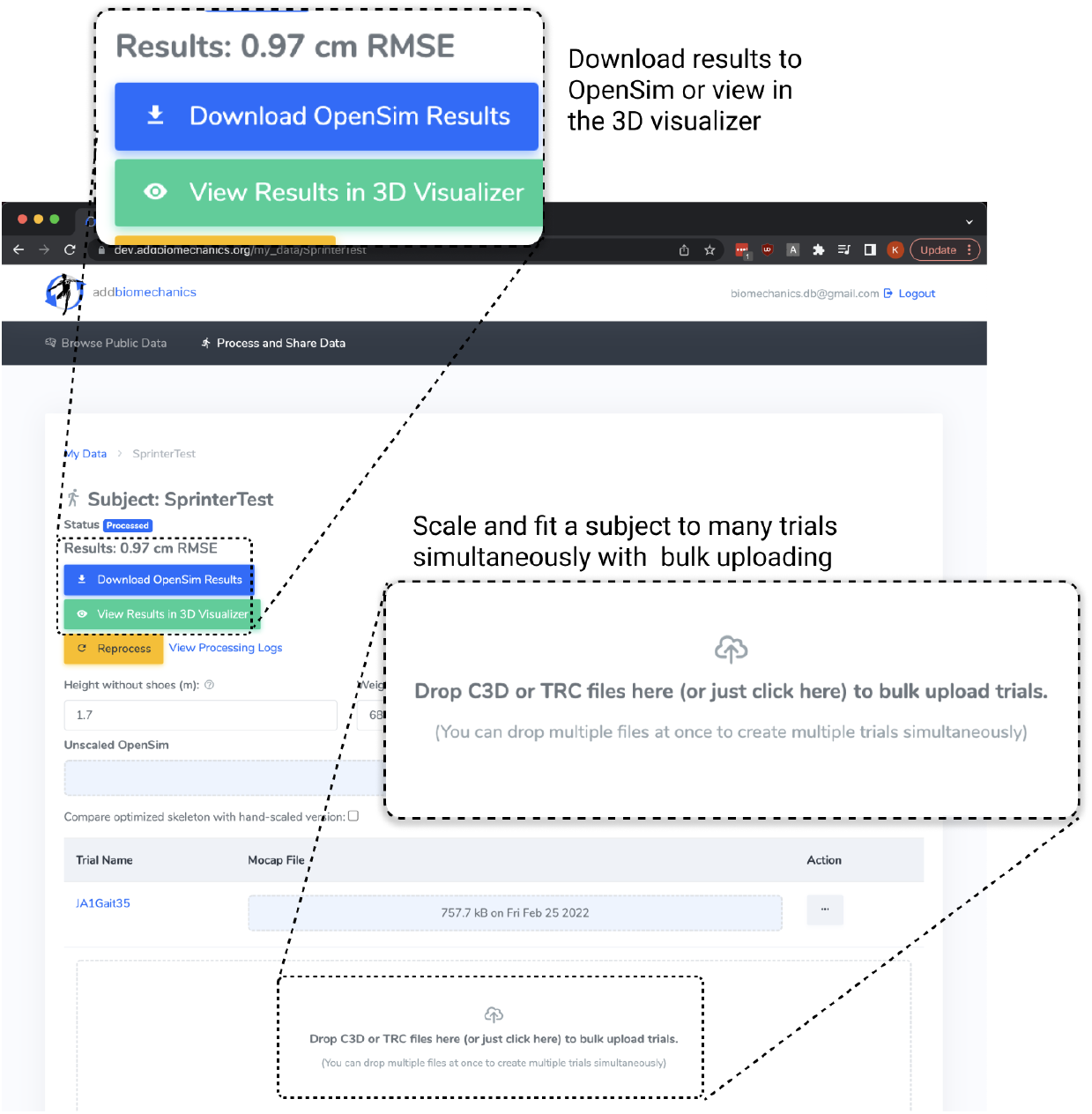
Screenshot of the AddBiomechanics user interface. The web interface allows users to drag and drop trials, and get subjects processed in the cloud within minutes.

Our open source implementation can optimize any user-defined OpenSim model with any marker set, and can operate on any number of marker trials per subject. The implementation finds three optimal scaling parameters **s_i_** = (*X, Y, Z*) for the *i*’th body segment in the model, and the optimal marker registration in the attached body segment’s local coordinate frame ***p_j_*** = (*X, Y, Z*) for the *j*’th marker in the model. Our method assumes that each marker is assigned to a body segment by the user. These values are returned to the user in the form of a scaled and registered (referred to as “optimized”) version of the OpenSim model the user submitted to the tool. The implementation also returns the inverse kinematics solution on the optimized OpenSim model for each uploaded trial.

### Evaluation

To evaluate our method on experimental data, we recruited three human experts who had recently collected and processed experimental motion capture data as part of three separate research studies. Expert 1 collected data for a gait perturbation-recovery study using a marker set based on [56] and analyzed it using the model from [48], Expert 2 analyzed the sprinting data from [57] using the model from [49], and Expert 3 collected data for sit-to-stand, squatting, jumping, and walking using a custom marker set and analyzed it using the model from [49]. Each expert scaled and registered between 4 and 17 trials, for a total of 34 trials. We then used the AddBiomechanics software to automatically compute body segment scales, marker registration, and inverse kinematics from the same raw experimental marker data.

Quantitative comparison of the solved joint angles with the ground truth joint angles is another critical test of our method. However, ground truth joint angles cannot be directly measured from the real life lab experiments. As a result, we generated synthetic marker trajectory data from a subject model with known body segment scalings, marker offsets, and joint angle trajectories and then attempted to recover the known joint angle trajectories from the synthetic marker data.

To generate synthetic data, we used several scaled OpenSim models, registered marker sets, and corresponding joint angle trajectories selected from the data created by our experts. We calculated marker positions at each frame of motion, assuming all markers were completely determined by the model and marker set (e.g. no non-rigid effects). We then ran AddBiomechanics on the synthetic marker data, giving it the generic unscaled version of the corresponding OpenSim model, and an unregistered version of the appropriate marker set to optimize, and compared the recovered motion to the known joint angles.

## Results

Our bilevel optimization algorithm to find body segment scales, marker offsets, and joint angle trajectories reduced the average root-mean-square marker reconstruction error by approximately 40% compared to a group of experts manually processing the same motion capture data (Fig 4), and faithfully reconstructed known joint trajectories from synthetic motion data (Fig 5, 6). The automated approach required no expert intervention.

**Fig 4.**
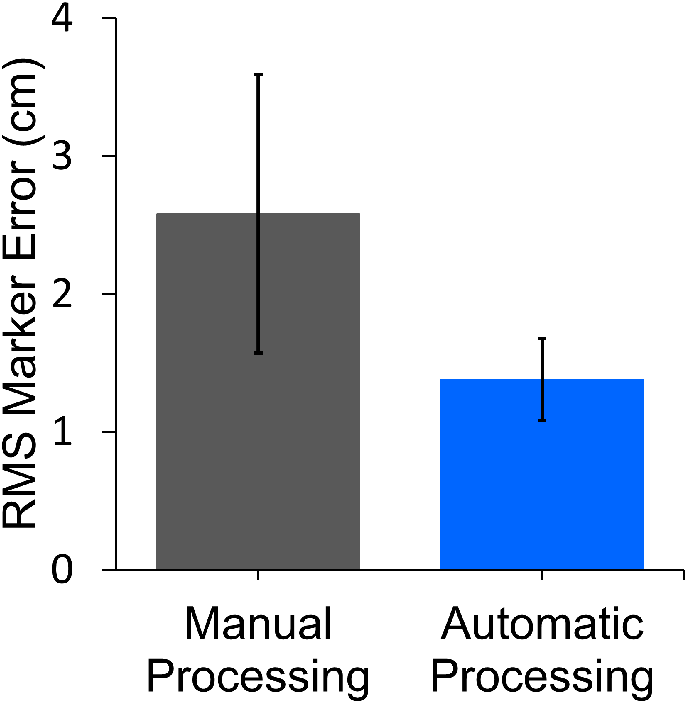
Manual Expert Processing vs. Automatic Processing. Automated scaling and marker fitting reduces the marker reconstruction error compared to manual expert processing. The bars show the average of per-trial marker RMSE, averaged over the 34 trials in the evaluation. The lines show the standard deviation of the marker RMSE across the trials, highlighting the difference in consistency between the automatic and manual processing.

**Fig 5.**
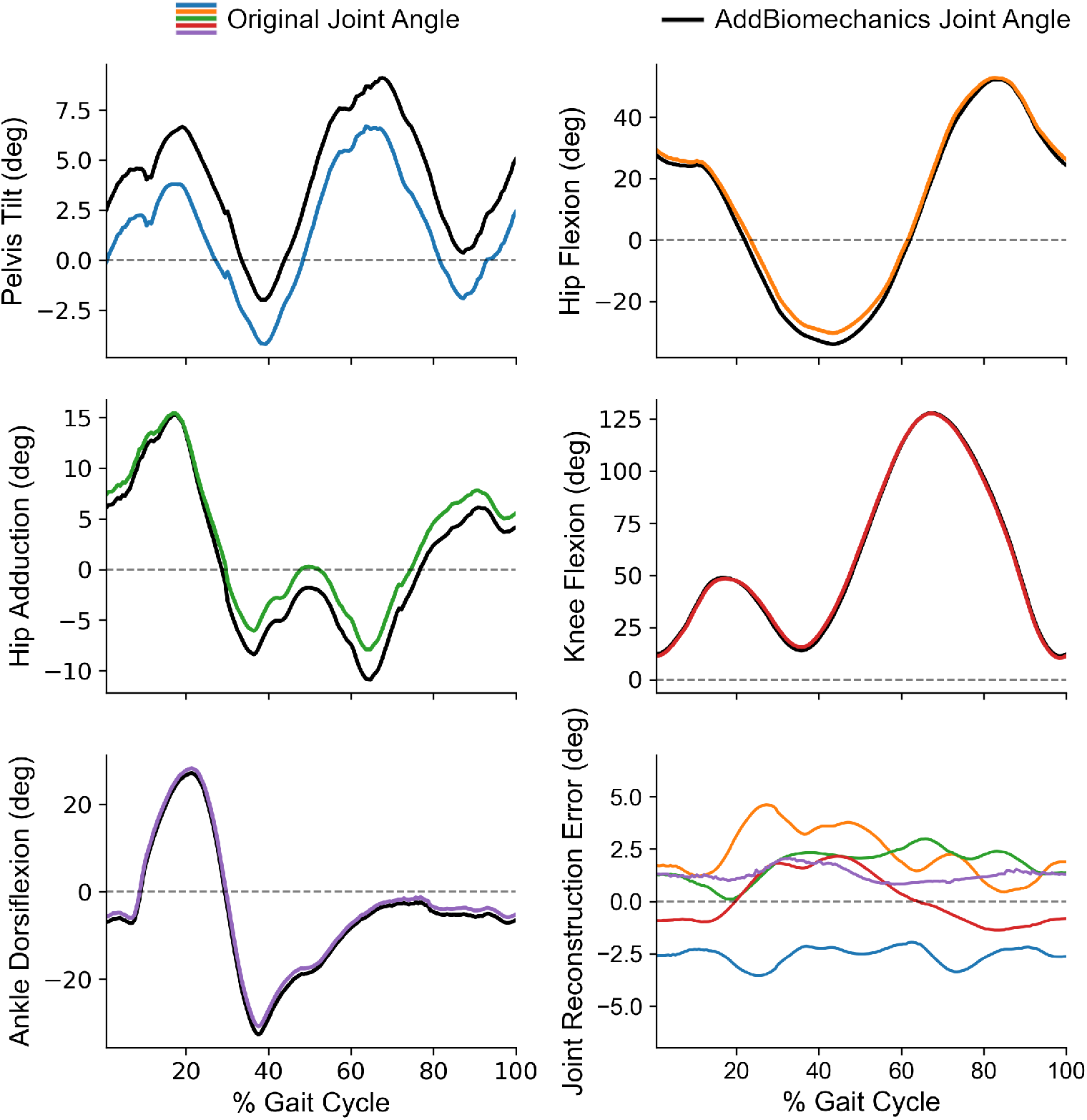
Comparison of original joint angles and AddBiomechanics joint angles on a sprinting trial.

**Fig 6.**
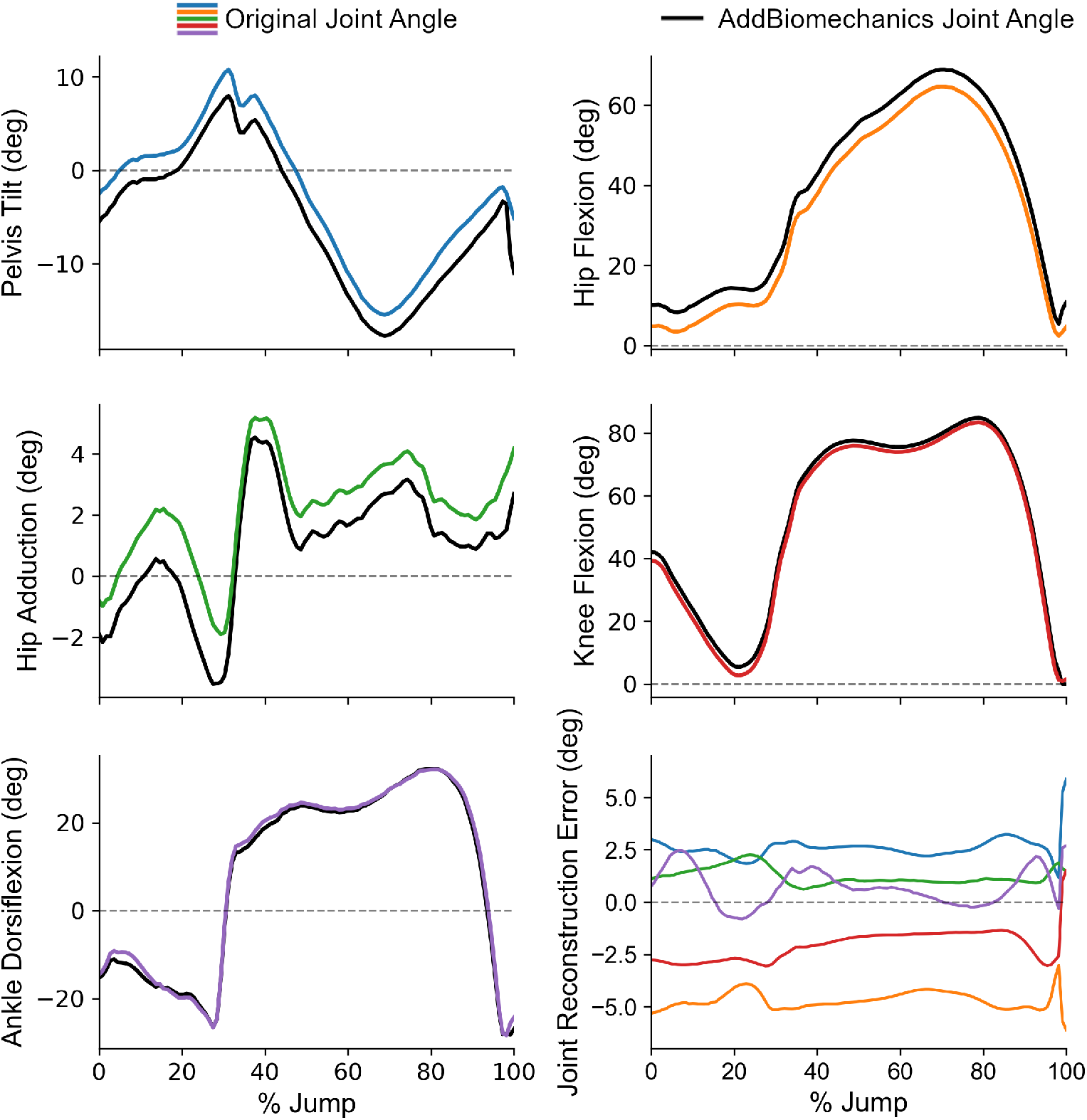
Comparison of original joint angles and AddBiomechanics joint angles on a jumping trial.

### Comparison to Human Experts Processing Experimental Data

The automated pipeline had significantly lower errors than manual processing by experts and a smaller standard deviation (Fig 4), where errors were measured as RMSE discrepancy between experimentally measured markers and simulated markers. This indicates that the automated system is able to produce combinations of scaling, marker registration, and inverse kinematics that more closely replicate experimental data.

Our experts perceived the manual body segment scaling and marker registration to be labor intensive: our three experts reported that manual processing required approximately one working day per subject. Average computation time for a trial processed with AddBiomechanics was 3 minutes on a consumer laptop.

Low marker reconstruction error is a necessary but not sufficient condition to establish the correctness of our system: it is possible to score well on marker RMSE and still have a model of the subject with hands, feet, and head that are scaled in an unrealistic way. This is possible because unrealistic bone scaling can be compensated for by adjusting marker registrations. We thus improved our confidence in the correctness of the automated scaling by fitting a multivariate gaussian over many individuals’ body measurements on the publicly available ANSUR II anthropometric dataset [50], and measuring how automated scaling compared with expert scaling on that gaussian. For each subject, we measured ***P***(automated)/P(expert), and found that across our comparison dataset the ratio was 1.65 ± 0.43. The value greater than 1 indicates that autoscaling consistently produced a more probable skeleton than expert scaling did.

### Comparison to Ground Truth Joint Angles from Synthetic Marker Data

We found that AddBiomechanics was able to recover the ground truth joint angles from synthetic marker data to an average of 0.79 deg RMSE and a maximum of 1.0 deg RSME (computed over all joints in a trial together) for motions including a standing calibration pose, walking, sprinting (Fig 5), and jumping (Fig 6). Representative joint angles are shown for sprinting and jumping, with the full results included as Table 1 in the supplementary data.

## Discussion

In addition to being computationally efficient, our method is able to recover joint angles on realistic synthetic data at state-of-the-art accuracy and outperform human experts in marker reconstruction error on experimental data. For comparison, the method in [24] assumed that all the body segment scalings were known to the algorithm and only attempted to find the marker offsets and the joint angles, and it reported 1.21 degree joint angle RMSE. Our method solves a harder problem, because it must recover segment scaling information from the data, and yet achieves similar results, with joint angle RMSE ranging from 0.46 degrees to 1.05 degrees. Our findings on marker reconstruction error on experimental data are also consistent with previous work on optimization based approaches, which all outperform human experts when fitting a model to the same data [24,44,45,58–60]. However, previous approaches required large amounts of compute time, were limited to one specific skeleton, or only addressed part of the body segment scaling and marker registration problem.

Our method goes from labeled marker trajectories to a scaled and registered musculoskeletal model and corresponding human motion in a few minutes on a standard consumer laptop or a low-end server. We also provide a web version where processing servers are hosted in the cloud, which means that end-users need no compute resources except a web browser. The new tool could open the possibility of movement biomechanics studies to a wider set of clinicians and researchers that do not possess the technical expertise or time traditionally required to achieve high-quality results. The manual processing for our small study of expert data took approximately a day of self-reported researcher time per subject, which was replaced by a few minutes of automated processing on an ordinary consumer laptop per subject. Even if the automated processing had only been comparable in quality, the labor savings alone would be a major improvement. Our comparison to manual expert processing also showed more accurate and consistent results. The wider significance of this method and the cloud-based tool will be to drive down cost and increase quality, consistency, and availability of biomechanical data analysis.

The optimizer does not always achieve a sufficiently accurate reconstruction of the joint angles for several reasons. First, there is some fundamental ambiguity in reconstructing so much information (body segment scales, marker offset registrations, and body positions) from only marker location data. For example, the pelvis can be tilted slightly forward, with the markers at the front of the pelvis shifted downward, and if the angles of the hips and spine are appropriately adjusted then the markers will still closely match the target data. Fig 6 provides an example of this happening on the jumping subject in our synthetic data experiments. If this effect is observed in practice, AddBiomechanics users can leverage the fact that the optimizer will prioritize solutions that move the markers as little as possible, and adjust the marker starting locations on the bones to more closely match the experimental placement. Secondly, the optimizer applies a statistical prior to body segment scales to bring them more in-line with population statistics as represented by the ANSUR II anthropometric dataset [50]. If the optimizer can find a way to fit the marker data with a skeleton that is more likely to exist in the ANSUR II population (such as by tilting the pelvis forward 2 degrees), it will choose that one, even if the “true” underlying skeleton was slightly different. The data in ANSUR II is large and detailed, but was collected from active duty military personnel, and so is not reflective of many patient populations. A broader anthropometric dataset could help address this limitation.

Our method also does not incorporate any dynamic quantities, ignoring ground reaction force data even when it is available. Ground reaction forces could provide very useful constraints on the acceleration of the center of mass of the subject, but exploiting these constraints to produce more accurate models is left to future work.

The open source cloud-based tool addbiomechanics.org reduces the barrier for researchers to integrate the optimization approach into their workflow. Researchers can get their data processed for free, if they agree to share the resulting anonymized motion data with the scientific community under a creative commons license. While large-scale public datasets of human motion already exist (e.g. [20]), they are focused on computer graphics applications and do not provide biomechanically accurate models of the experimental subjects. Because AddBiomechanics is focused on human motion biomechanics, the cloud application also encourages sharing ground reaction force data along with kinematics, even though the optimizer does not currently leverage this data to increase the accuracy of its solutions. We hope AddBiomechanics will enable creation of a large-scale public dataset of accurately modeled human motion biomechanics, including both state-of-the-art models of experimental subjects and measurements of biomechanically relevant quantities such as ground reaction forces, electromyography, and others, as the tool grows in the future.

## Data Availability Statement

All data and code used for running experiments, model fitting, and our cloud application is available on a GitHub repository at https://github.com/keenon/AddBiomechanics and we have archived our code on Zenodo (DOI: 10.5281/zenodo.6981568).

## Acknowledgments

This work was supported by NSF GRFP (DGE-1656518), The National Institutes of Health (P41-EB027060 and P2C-HD101913), the Hoffman-Yee Research Grants (Stanford-HAI-203112), Stanford Bio-X SIGF Paul Berg Interdisciplinary Biomedical Graduate Fellowship, the Wu Tsai Human Performance Alliance and Joe and Clara Tsai Foundation. Special thanks to Carmichael Ong, who lent us his OpenSim expertise and helped create several skeletons and markersets for AddBiomechanics, and Reed Gurchiek and Nick Bianco for early feedback on the tool.

## Supplementary

**Table 1.** Synthetic Data Joint Angle Recovery.

| Source | Motion Type | Model | RMSE Joint Error (deg) |
| --- | --- | --- | --- |
| Expert 1 | Calibration | Rajagopal [48] | 1.05 deg |
| Expert 1 | Walking | Rajagopal [48] | 0.97 deg |
| Expert 2 | Sprinting | Lai Arnold [49] | 0.46 deg |
| Expert 3 | Jumping | Lai Arnold [49] | 0.68 deg |

The underlying ground truth is recovered with approximately 1 degree of joint angle error across a range of motion types, despite fundamental ambiguity.

## Notes

### Competing Interest Statement

The authors have declared no competing interest.

